# Elongated Hypocotyl 5 (HY5) regulates *BRUTUS* (*BTS*) to maintain Iron homeostasis in *Arabidopsis thaliana*

**DOI:** 10.1101/2022.04.26.489524

**Authors:** Samriti Mankotia, Dhriti Singh, Kumari Monika, Himani Meena, Varsha Meena, Ram Kishor Yadav, Ajay Kumar Pandey, Santosh B. Satbhai

## Abstract

Iron (Fe) is an essential micronutrient for both plants and animals. Fe limitation significantly reduces crop yield and therefore has adverse impacts on human nutrition. Owing to limited bioavailability of Fe, plants have adapted different strategies that regulate Fe uptake and homeostasis. Particularly, modifications of root growth traits are a key for the survival of plants on Fe-deficient soils. Understanding the molecular basis for these root growth responses will have critical implications for plant breeding. Fe uptake is regulated by a cascade of basic helix-loop-helix (bHLH) transcription factors. In our study, we show that *HY5* (Elongated Hypocotyl 5), a member of the basic leucine zipper (bZIP) family of transcription factors plays an important role in the Fe deficiency signalling pathway in *Arabidopsis thaliana*. The *hy5* mutant plants failed to mount an optimum Fe deficiency response and showed severe root growth defects under Fe limitation that could be partially reverted by complementation of *hy5* mutant. qRT-PCR analysis revealed that the induction of the genes involved in Fe uptake pathway [FER-like iron deficiency-induced transcription factor (*FIT*), Ferric Reduction Oxidase 2 (*FRO2*) and Iron-Regulated Transporter1 (*IRT1*)] is significantly reduced in the *hy5* mutants as compared to the wild-type plants under Fe deficiency. Moreover, we also found that HY5 function is critical for activating the expression of coumarin biosynthesis genes (*F6’H1, S8H, PDR9* and *BGLU42)* under Fe deficiency. Interestingly, our results showed that *HY5* acts as a negative regulator of BRUTUS (*BTS)* which is known to negatively regulate Fe deficiency response. Chromatin Immunoprecipitation followed by qPCR revealed direct binding of HY5 to the promoter of *BTS*. Altogether, our results showed that *HY5* plays an important role in regulation of Fe deficiency responses in Arabidopsis.

## INTRODUCTION

Iron (Fe) is an indispensable micronutrient for plant growth and development. It plays an irreplaceable role in many crucial processes like respiration, photosynthesis, hormone biosynthesis, pathogen defense and chlorophyll biosynthesis (Hänsch and Mendel, 2009; Balk and Schaedler, 2014). Plants mainly obtain Fe from the soil. Although Fe is present in abundance yet most of it is unavailable for absorption by plants. At basic and neutral pH in the well-aerated soils, it tends to form insoluble crystals of ferric (Fe^3+^) oxyhydrates, which cannot be taken up by the plants (Thomine and Lanquar, 2011)

For efficient Fe uptake from the soil, plants have adopted two types of strategies: reduction based strategy, also known as strategy I and chelation based strategy, also known as strategy II. *Arabidopsis thaliana* utilizes reduction-based strategy (Strategy I) for Fe uptake from soil (Kobayashi and Nishizawa, 2012; Brumbarova, Bauer and Ivanov, 2015). In this strategy, protons are released into the rhizosphere from the root PLASMA MEMBRANE PROTON ATPASE 2 (AHA2), which acidifies the rhizosphere, increasing Fe solubility in soil. In addition to this, Fe^3+^ is also chelated and mobilized by coumarin family phenolics released into the rhizosphere by the PDR9 (ABCG transporter) (Td, Hsuan and Schmidt, 2017). The resulting solubilized Fe^3+^ is reduced to Fe^2+^ by the FERRIC REDUCTION OXIDASE 2 (FRO2) enzyme (Connolly *et al*., 2003). The Fe^2+^ is taken inside the root epidermal cells by the IRON-REGULATEDTRANSPORTER1 (IRT1) transporter (Vert *et al*., 2002; Santi and Schmidt, 2009; Fourcroy *et al*., 2016).

The expression of Fe uptake genes is induced under Fe deficiency so as to absorb more Fe from the rhizosphere. FIT (Fer like Iron deficiency Transcription Factor) heterodimerize with subgroup Ib bHLH TFs (bHLH38, bHLH39, bHLH100 and bHLH101) to activate the expression of *FRO2* and *IRT1* (Yuan *et al*., 2008; Wang *et al*., 2013; Liang *et al*., 2017). The expression of subgroup Ib bHLH TFs is also induced under Fe deficiency and their expression is activated by subgroup IVc bHLH TFs (bHLH34, bHLH104, bHLH105 (ILR3), and bHLH115 (Wang *et al*., 2013). ILR3, in addition to its role as a transcriptional activator also acts as transcriptional repressor by interacting with PYE and negatively regulate the expression of genes involved in Fe transport (*NAS4*), storage (*AtFER1, AtFER3, AtFER4, VTL2*) and assimilation (*AtNEET*) (Tissot *et al*., 2019). The subgroup IVc bHLH transcription factors are expressed under both Fe sufficient and Fe deficient conditions which indicates that they are regulated at protein level so that they can induce the expression of subgroup Ib genes only under Fe-deficient conditions. The BRUTUS (BTS) which is a E3 ubiquitin ligase is involved in regulating the subgroup IVc bHLH TFs at protein level (Long *et al*., 2010; Selote *et al*., 2015). It degrades subgroup IVc bHLH TFs under Fe sufficient conditions negatively regulating Fe homeostasis to avoid Fe overload (Selote *et al*., 2015). The BRUTUS LIKE1 (BTSL1) and BTSL2 which are two closely related RING E3 ubiquitin ligases also negatively regulate Fe homeostasis by directly targeting FIT for degradation by the 26S proteasome (Rodríguez-Celma *et al*., 2019). IRON MAN peptides (IMAs), a class of small peptides induced under Fe deficiency have been recently shown to play a role in activating Fe deficiency response by sequestering BTS and promoting accumulation of bHLH105 and bHLH115 (Grillet *et al*., 2018; Li *et al*., 2021). bHLH121 (URI) has also been found to positively regulate Fe homeostasis by interacting with bHLH IVc TFs (Kim *et al*., 2019; Gao *et al*., 2020; Lei *et al*., 2020).

HY5 (Elongated Hypocotyl 5) is a member of the basic leucine zipper family of transcription factors (bZIP) and crucial regulator of photomorphogenesis. *hy5* mutants have less chlorophyll content, elongated hypocotyl and increased number of lateral roots as compared to the wild type plants (Oyama, Shimura and Okada, 1997b; Ang *et al*., 1998; Holm *et al*., 2002). It is also known to positively regulate, copper and sulfur signalling pathways (Jonassen, Lea and Lillo, 2008; Lillo, 2008; Lee, Koprivova and Kopriva, 2011; Yanagisawa, 2014; Zhang *et al*., 2014; Huang *et al*., 2015). There are some reports which indicate *HY5* to have a broader role in Fe signalling. Based on gene coexpression analysis, *HY5* is speculated to be a master regulator of Fe homeostasis (Brumbarova and Ivanov, 2019). Many Fe-responsive genes contain HY5 binding sites in their promoters in *Arabidopsis* (Vélez-Bermúdez and Schmidt, 2022). Recently, in tomato *HY5* has been shown to be involved in regulating Fe uptake. The phyB gets activated by red light and induces HY5 which then moves from shoot to roots and triggers Fe uptake by inducing *FER* expression (Guo *et al*., 2021). However, the role of HY5 in Fe signalling has not been studied in detail till now and need to be investigated.

In this study, we report that *HY5* is involved in regulating Fe deficiency responses in *Arabidopsis. HY5* function is crucial for primary root growth under Fe limiting conditions. We found that HY5 is required for regulating the expression of genes involved in maintaining Fe homeostasis which include *IRT1, FIT, FRO2, PYE* and *BTS* as well as genes involved in coumarin biosynthesis *CYP82C4, S8H, F6’H1* and *BGLU42*. Our data also indicate that HY5 acts as a negative regulator of BRUTUS (*BTS)* to modulate Fe deficiency responses in Arabidopsis. We conclude that HY5 acts as a key player to regulate genes involved in Fe homeostasis pathway in Arabidopsis.

## RESULTS

### *hy5* mutants have an impaired Fe-deficiency response

To investigate the role of *HY5* in Fe homeostasis, we used a T-DNA insertional mutant of *hy5* and compared its phenotype with the wild-type (WT) plants. For phenotypic analysis, both WT Col-0 and *hy5* mutants were grown on half-strength Murashige and Skoog (MS) medium containing Fe (+Fe) and half-strength MS medium lacking Fe (-Fe). It is known that the primary root length of WT plants increases under moderate Fe deficiency and decreases under severe Fe deficiency (Gruber *et al*., 2013). Our phenotypic analysis revealed that there was a significant increase in the primary root length of the WT seedlings when grown on -Fe as compared to the +Fe but in case of the *hy5* mutant seedlings root length significantly decreased when grown on -Fe as compared to +Fe (Figure 1A, B). The *hy5* mutants were stunted and leaves were more chlorotic as compared to the WT plants which indicates that *hy5* mutant has less chlorophyll content than WT under Fe-limiting conditions. To confirm this, total chlorophyll content of both WT and *hy5* mutants grown on +Fe and -Fe media for 10 days was quantified and it was found that under +Fe conditions both *hy5* and WT has similar chlorophyll content but under -Fe conditions *hy5* mutants have significantly lower chlorophyll content as compared to WT plants (Figure 1C).

**Figure 1.**
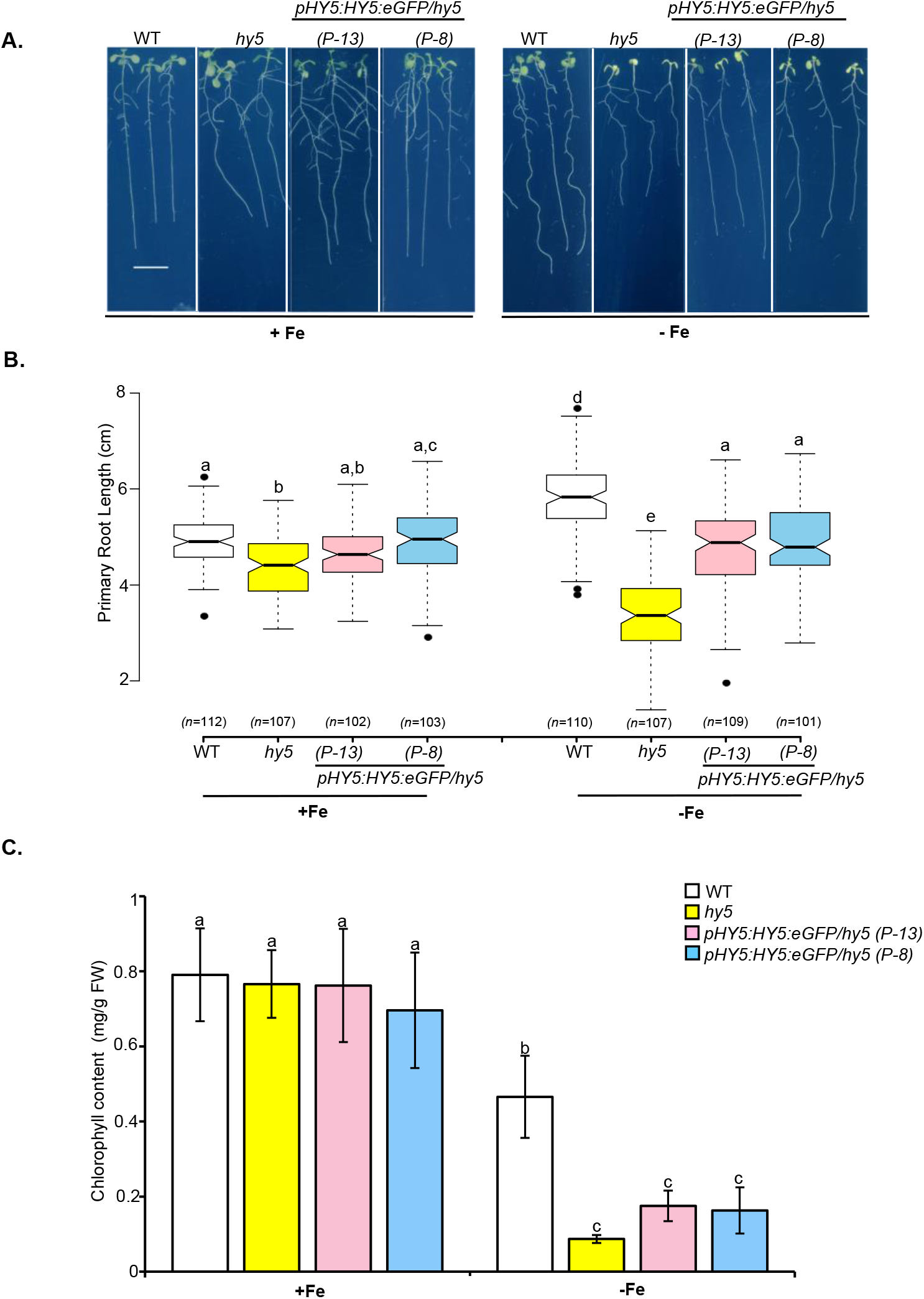
*hy5* mutants have less tolerance to iron deficiency. **A**. Phenotypes of the *Arabidopsis* wild type (WT), *hy5* and *pHY5:HY5:eGFP/hy5* (*P-13* and *P-8*) grown for 10 days on Fe-sufficient and Fe-deficient medium. Scale bars, 1 cm. **B**. Boxplot of root length of the wild type (WT), *hy5* and *pHY5:HY5:eGFP/hy5* (*P-13* and *P-8*) grown for 10 days on Fe-sufficient and Fe-deficient medium. **C**. Chlorophyll content of the wild type (WT), *hy5* and *pHY5:HY5:eGFP/hy5* (*P-13* and *P-8)* grown for 10 days on Fe-sufficient and Fe-deficient medium. Data shown is an average of three independent experiments. Each experiment consists of 4 biological replicates and each replicate consists of a pool of around six seedlings. Error bars represent ±SEM. Means within each condition with the same letter are not significantly different according to one-way ANOVA followed by post hoc Tukey test, P < 0.05.

To confirm that these phenotypes are because of loss of *HY5* function, we generated complementation lines by transforming the *HY5* translational fusion construct (*ProHY5:HY5:eGFP*) into *hy5* mutant to restore its function. We then selected two independent T3 complementation transgenic plants P-13 and P-8 (*ProHY5:HY5:eGFP/hy5*) and to confirm the lines we checked the HY5 protein expression in the *ProHY5:HY5:eGFP/hy5* (P-13 and P-8) complementation lines using confocal microscopy. We found that HY5 is uniformly expressed in all the cell layers in root (Figure S1). The root length defects were partially rescued in both the complementation lines (Figure 1A, B). Chlorophyll content in *hy5* mutant and complemented lines was comparable to WT plants under +Fe conditions but under -Fe conditions as compared to WT, chlorophyll content in *hy5* mutants was strongly reduced (80.89%), however in both the complemented lines the chlorophyll content reduction was suppressed (62.37% and 63.15%) as compared to *hy5* mutant plants (Figure 1C). This results also indicates that the chlorophyll content reduction defects of *hy5* mutants was slightly rescued in both the complemented lines under -Fe conditions. The observed partial rescue phenotypes might be due to lower levels of HY5 protein in complemented lines as compared to WT plants.

*HYH* (*HY5* homolog) is known to act redundantly with *HY5* in regulation of nitrogen signalling, inhibition of hypocotyl growth and lateral root development, and expression of light-inducible genes (Oyama, Shimura and Okada, 1997a; Ang *et al*., 1998; Holm *et al*., 2002; Gangappa and Botto, 2016). So, in to order to find out whether it has any role in Fe deficiency, we used *hyh* mutant line and compared the phenotype of *hyh* mutants with the *hy5* mutant, WS (Wassilewskija) and *hy5/hyh* double mutants under -Fe conditions. The *hyh* mutants showed similar primary root length to wild-type (WS) plants, however *hy5* single mutants displayed a significant inhibition of primary root growth under -Fe (Figure S2). This result obtained using *hy5* mutants from WS background independently confirms root growth phenotype of *hy5* mutants from Col-0 background (Figure 1) under -Fe. Moreover, the root length of *hy5/hyh* double mutant plants was comparable with *hy5* single mutant plants. These results clearly indicate that *HY5* works independently of *HYH* under -Fe conditions and *HYH* function is not critical under -Fe condition.

Next, we checked whether *hy5* mutants show Fe-deficiency specific growth responses in soil. For this, WT, *hy5* and *ProHY5:HY5:eGFP/hy5* (P-13 and P-8) were grown on control soil (pH 5.0 to 6.0) and alkaline soil (pH 7.0 to 8.0) which creates Fe-limiting conditions because Fe solubility decreases with increase in pH. Alkaline soil was made by adding calcium oxide in the control soil. It was found that under alkaline soil conditions, *hy5* mutants were more chlorotic as compared to the WT (Figure 2A). Moreover, we also observed that the chlorotic phenotype of *hy5* mutant plants was partially rescued in the *ProHY5:HY5:eGFP/hy5 (P-13 and P-8)* lines (Figure 2A). To confirm this phenotype is specifically due to Fe limitation at higher pH, we added the external Fe (Fe-EDDHA) to alkaline soil and compared the phenotype with the plants grown under control and alkaline soil conditions. When external Fe was provided, we found that the chlorotic phenotype of *hy5* mutants was rescued (Figure 2A). Altogether, these results indicate that *hy5* mutant is sensitive to -Fe conditions as compared to the WT.

**Figure 2.**
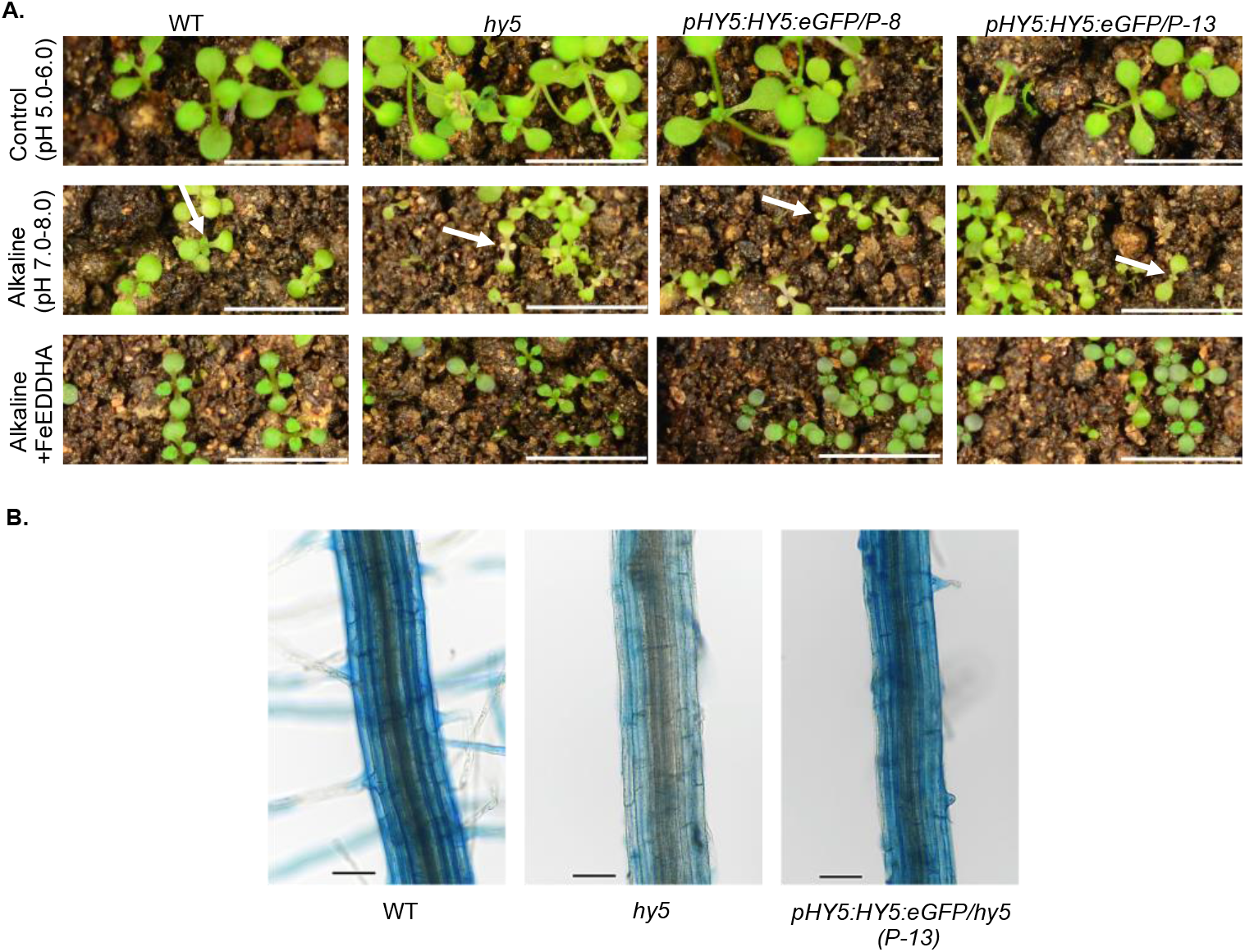
*hy5* mutant is more sensitive to iron-deficiency and have less iron content as compared to WT. **A**. Phenotypes of the WT, *hy5, pHY5:HY5:eGFP/hy5 (P-13 and P-8)* grown on normal soil (pH 5.0-6.0), alkaline soil (pH 7.0-8.0) and alkaline soil watered with Fe-EDDHA (Ethylenediamine di-2-hydroxyphenyl acetate ferric) for two weeks. Bars =10mm. **B**. Perl’s stained maturation zone of WT, *hy5* and *pHY5:HY5:eGFP/hy5* (P-13). Bars = 1mm. White arrows indicate true leaves undergoing chlorosis.

We also performed perl’s blue staining to see the impact of *HY5* mutation on Fe content and found that *hy5* mutant has less Fe content than WT. In the complemented line *ProHY5:HY5:eGFP/hy5 (P-13)*, the Fe content looks comparable to WT (Figure 2B) which suggests that Fe uptake is affected in the *hy5* mutants and HY5 function is important for optimum Fe uptake under -Fe conditions.

### HY5 regulates the expression of Fe-Deficiency-Responsive genes

Since *hy5* mutants have reduced tolerance to Fe-deficiency, we wanted to determine whether the expression of Fe-deficiency-responsive genes was also affected in the *hy5* mutants. We performed qRT-PCR to check the expression of genes involved in Fe-deficiency. This was done by examining the mRNA level of the major genes involved in controlling Fe homeostasis in WT and *hy5* seedlings grown on 1/2 MS for six days and then transferred to +Fe or –Fe (+300 µM Ferrozine) for three days. The expression of major Fe uptake genes (*IRT1, FRO2*) was found to be significantly reduced in the *hy5* mutant as compared to the WT plants under Fe deficiency (Figure 3A). Fe deficiency induced *FIT* expression was also found to be compromised in the *hy5* mutants as compared to the WT plants under Fe deficiency (Figure 3A). This indicates that *HY5* function is important for the induction of key genes involved in Fe uptake. The expression of *BTSL1, BTSL2, IMA1, IMA2, IMA3* and subgroup Ib bHLH TFs (*bHLH38, bHLH39, bHLH100 and bHLH101*) was not affected significantly in the mutant compared with the wild type (Figure S3 and Figure 3B).

**Figure 3.**
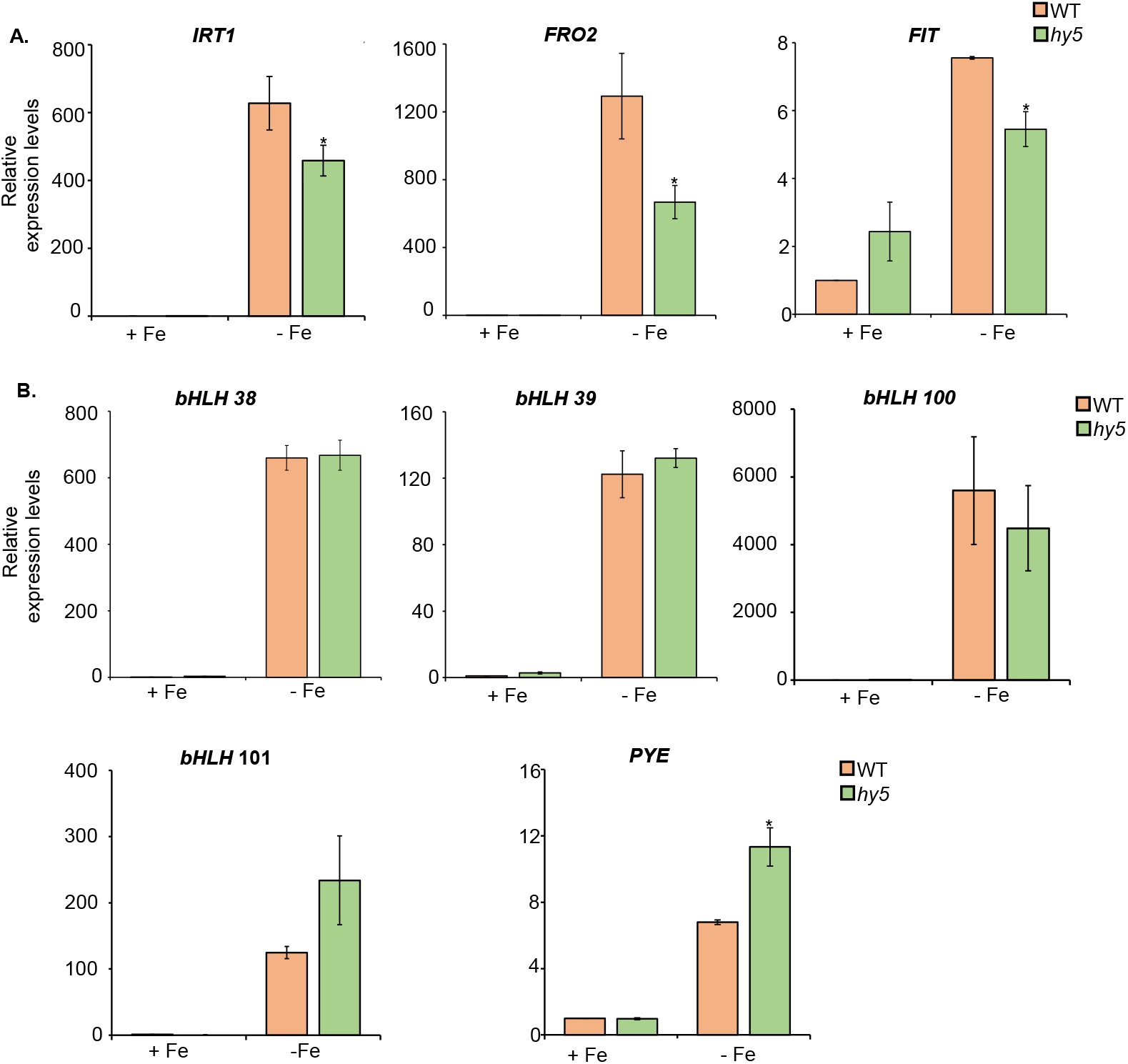
Expression of genes involved in iron homeostasis is affected in the *hy5* mutant. A. Expression levels of *IRT1, FRO2* and *FIT*. B. Expression levels of subgroup Ib bHLH TFs (*bHLH 38, 39,100* and *101*) and *PYE*. Relative expression was determined by qRT-PCR in WT and *hy5* mutant seedlings grown on +Fe media for 6 days and transferred to both +Fe and –Fe (+300µM Fz) for three days. Data shown is an average of three biological replicates (n=2 technical replicates). Each biological replicate consists of pooled RNA extracted from roots of ∼ 90 seedlings. Error bars represent ±SEM.*Significant difference by Student’s t test (P≤0.05).

We also checked the expression level of PYE, a TF which plays a negative role in the Fe deficiency responses (Long *et al*., 2010). The expression level of *PYE* which is known to negatively regulate genes involved in iron transport (*NAS4*), iron storage (*FER1, FER3, FER4* and *VTL2*) and iron assimilation (*NEET*) was found to be significantly more induced in the *hy5* mutant as compared to the WT (Figure 3B). This indicates that *HY5* negatively regulates *PYE* expression.

The qRT-PCR results indicate that there was reduced induction of *IRT1* expression in the *hy5* mutant as compared to WT under –Fe conditions. To further understand how *hy5* mutation affects the expression pattern of *IRT1* in roots we used *ProIRT1:GUS* line. We compared *ProIRT1:GUS* activity in the wild type and *hy5* background grown on both +Fe and –Fe for 8 days. In the wild type plants, we observed a strong induction of the reporter gene activity in plants grown on –Fe media but in case of the *hy5* mutant plants, we observed significantly lower induction of reporter gene activity as compared to WT plants (Figure 4). These results clearly indicate that *HY5* is important for optimum induction of *IRT1* under –Fe conditions.

**Figure 4.**
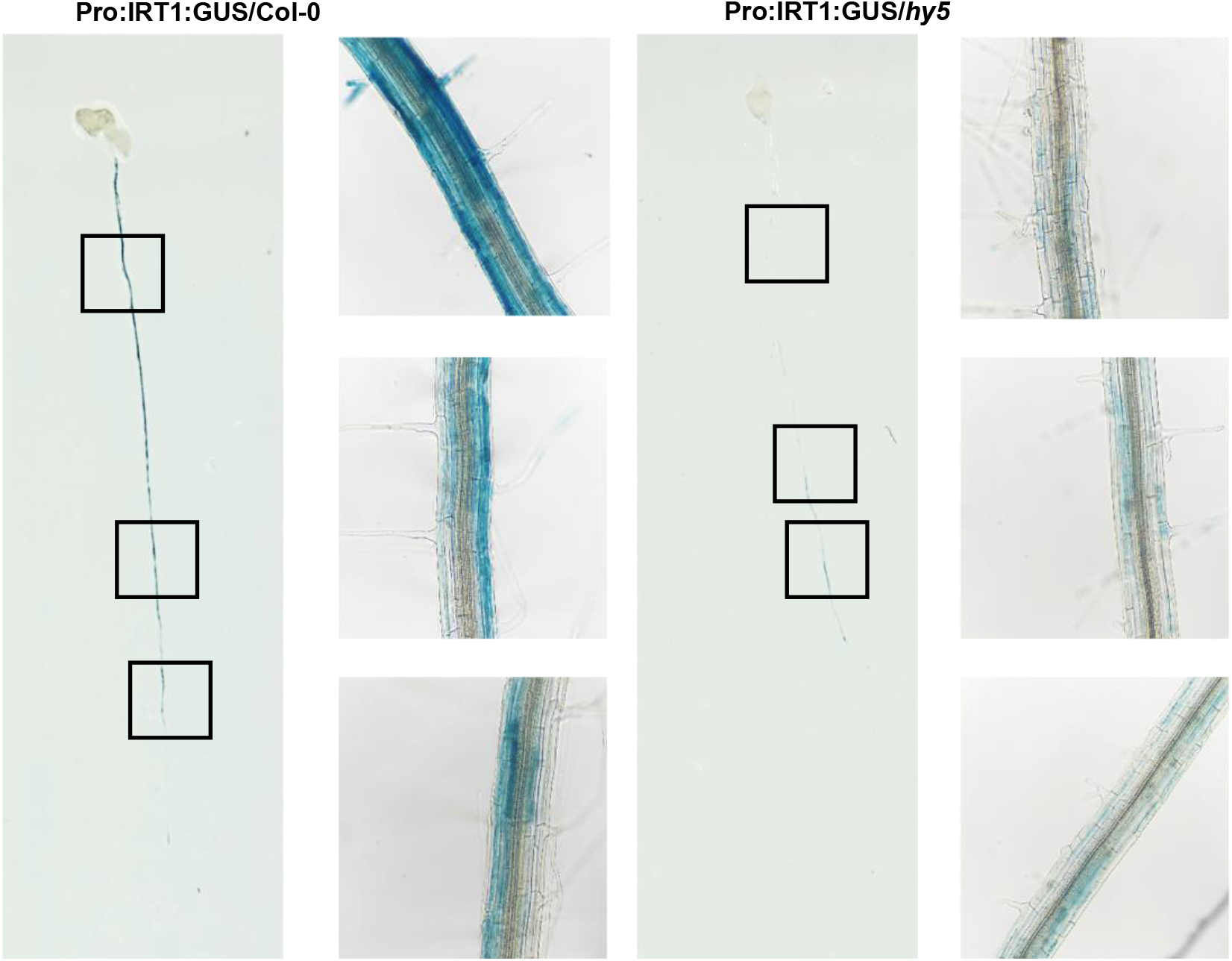
*IRT1* induction is compromised in the *hy5* mutant under -Fe. Analysis of *ProIRT1:GUS* gene activity in WT Col-0 and *hy5* mutant seedlings expressing it by GUS Staining. Plants grown under –Fe media for 8 days were stained for GUS activity.

Since *FRO2* induction was significantly lower in the *hy5* mutant as compared to the wild type. So, we checked ferric-chelate reductase (FCR) activity in the mutant and FCR activity was found to be significantly less induced in the mutant as compared to the wild type under -Fe (Figure S4). Collectively, these results indicate that *HY5* function is crucial for the activation of genes involved in Fe uptake under -Fe.

### HY5 functions upstream to BTS

BRUTUS (BTS), acts as a negative regulator of Fe uptake by controlling bHLH subgroup IVc protein levels through proteasomal degradation (Selote *et al*., 2015). Our qRT-PCR analysis clearly showed that *BTS* is induced more in the *hy5* mutant as compared to the WT under -Fe (Figure 5A). To determine whether *HY5* directly binds to promoter of *BTS* we analysed the promoter sequence of *BTS*. We found HY5 binding motifs, three G-boxes, one CG-Hybrid which are within 406 to 488 bp upstream (region I) and one CG-Hybrid which is 1400 bp upstream (region II) to the translational start site (ATG) in the promoter of *BTS* (Figure 5B). We hypothesized that HY5 directly binds on the *BTS* promoter to regulate its expression and performed ChIP-qPCR experiments to confirm the binding. ChIP experiments were performed using *hy5* mutant lines carrying the transgene *Pro:HY5:HY5:YFP* using an anti-GFP antibody. As a control, the same experiment was conducted on *35S:*e*GFP* seedlings. We found significant enrichment of *BTS* promoter region I with HY5 protein in *Pro:HY5:HY5:YFP/hy5* as compared to *35S:eGFP* (Figure 5C). We used IgG Ab as a negative control. This observation indicates that HY5 directly binds in vivo to the promoter of *BTS*.

**Figure 5.**
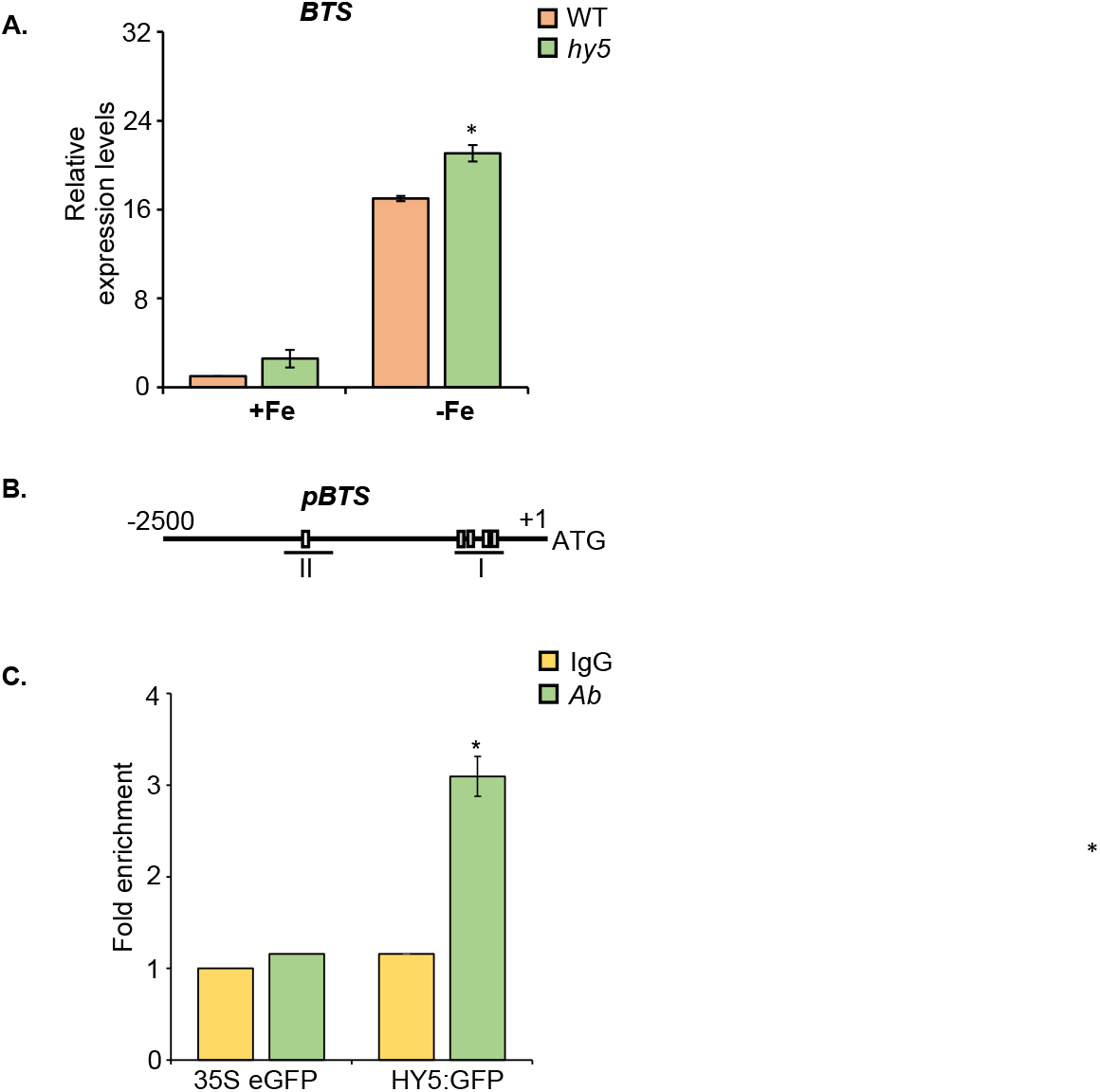
HY5 directly binds on *BTS* promoter. A. Expression levels of *BTS*. Relative expression was determined by qRT-PCR in WT and *hy5* mutant seedlings grown on +Fe media for 6 days and transferred to both +Fe and –Fe (+300µM Fz) for three days. Data shown is an average of three biological replicates (n=2 technical replicates). Each biological replicate consists of pooled RNA extracted from ∼ 90 seedlings. Error bars represent ±SEM.*Significant difference by Student’s t test (P≤0.05). B. Schematic diagram of the promoter of the *BTS*. White boxes represent CG-Hybrid, grey boxes represent G-boxes. Lines under the boxes represent sequences detected by ChIP qPCR. C. ChIP-qPCR showing relative enrichment of the *BTS* regulatory region I bound by HY5. The ChIP assay was performed using *pHY5::HY5:YFP/hy5* and *35S:eGFP* seedlings grown on +Fe media for 10 days. Error bars represent ±SEM..

To further investigate how *hy5* mutation affects the expression pattern of *BTS in planta*, we used *ProBTS:GUS* line. We compared *ProBTS:GUS* activity in the wild type and *hy5* background grown on both +Fe and –Fe for 8 days. In the *hy5* mutant plants, we observed a stronger induction of the reporter gene as compared to the wild type plants in which we observed significantly lower induction of reporter gene activity under -Fe conditions further confirming the results of qRT (Figure 6). These results clearly indicate that *HY5* negatively regulates *BTS* induction under –Fe conditions.

**Figure 6.**
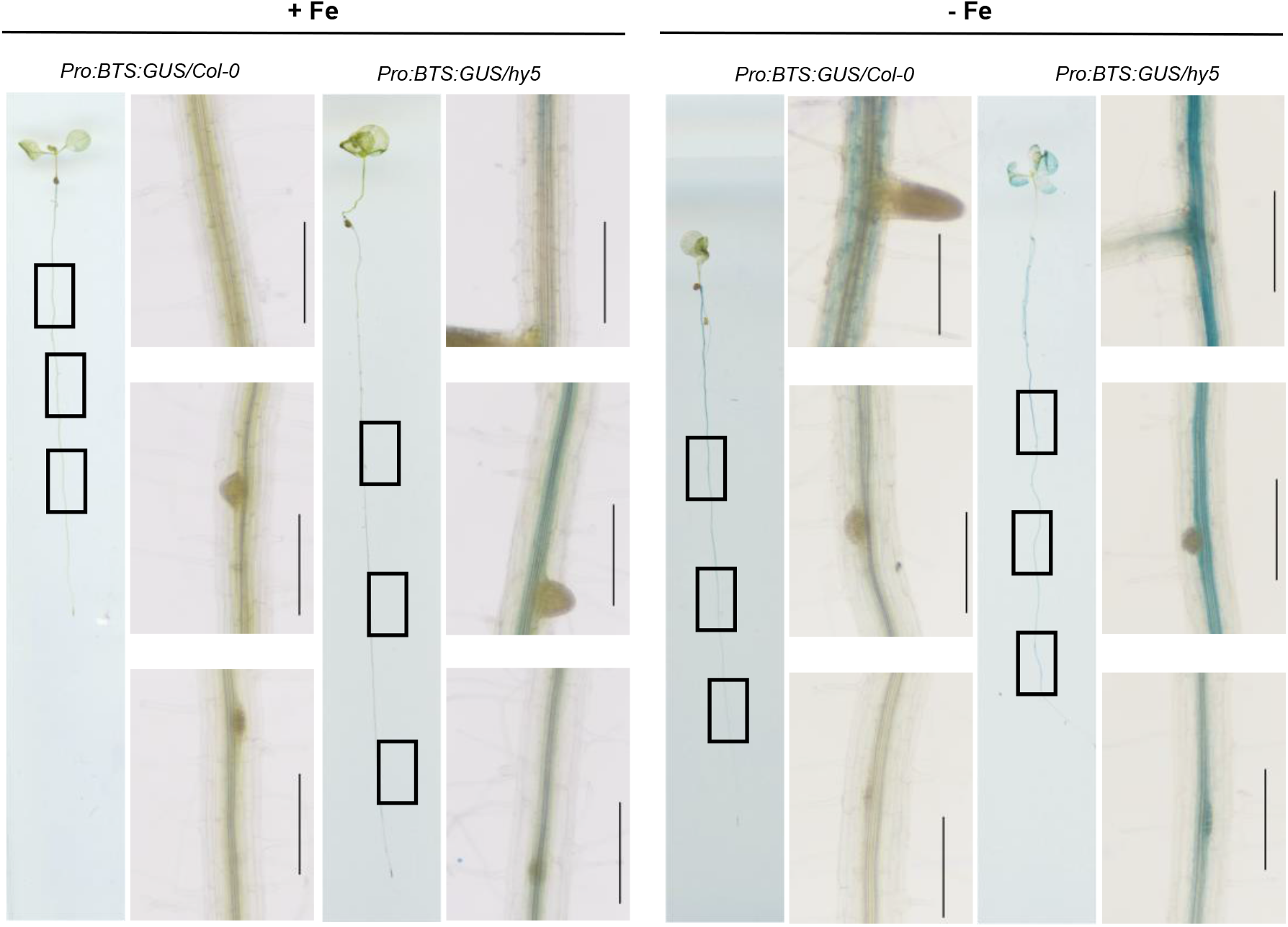
*BTS* is induced more in the *hy5* mutant. Analysis of *ProBTS:GUS* gene activity in WT Col-0 and *hy5* mutant seedlings. Plants grown under +Fe and –Fe media for 8 days were stained for GUS activity.

As we found that *BTS* expression is induced more in the *hy5* mutant as compared to the wild type and HY5 directly binds to its promoter. We hypothesised that *BTS* is a direct downstream target of *HY5*. To confirm this hypothesis, we generated *hy5/bts* double mutant by crossing *bts* and *hy5* mutant plants. Under control (+Fe) growth conditions, we did not observe significant difference in the primary root length between the *hy5/bts* double and the single mutants but under –Fe conditions, the *hy5/bts* double mutants showed phenotype similar to *bts* mutants and did not show decrease in primary root length as *hy5* mutants. (Figure 7A, B). We also checked the phenotype on alkaline soil and we found that on alkaline soil the leaves of *hy5/bts* double mutant were not chlorotic like *hy5* mutant (Figure S5). These results indicate that *bts* mutation suppresses the *hy5* mutant phenotype or *BTS* is epistatic to *HY5*.

**Figure 7.**
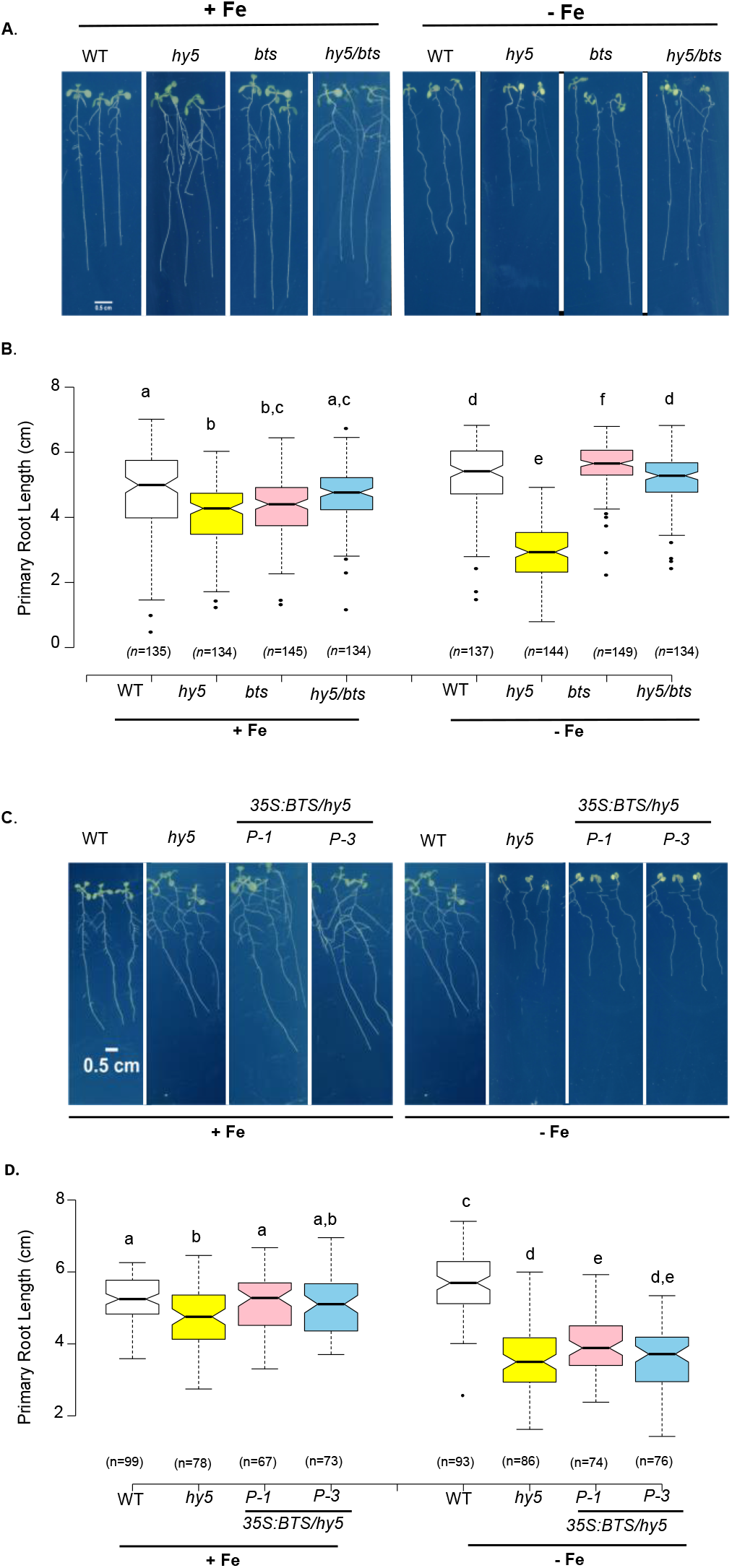
*HY5* acts upstream to BTS. A. Phenotypes of the *Arabidopsis* wild type (WT), *hy5, bts* and *hy5bts* grown for 10 days on Fe-sufficient and Fe-deficient medium. Scale bars, 0.5 cm. B. Boxplot of root length of the wild type (WT), *hy5, bts* and *hy5bts* grown for 10 days on Fe-sufficient and Fe-deficient medium. C. Phenotypes of the *Arabidopsis* wild type (WT), *hy5* and *35S:BTS/hy5 (P-1 and P-3)* grown for 10 days on Fe-sufficient and Fe-deficient medium. D. Boxplot of root length of the wild type (WT), *hy5* and *35S:BTS/hy5 (P-1 and P-3)* grown for 10 days on Fe-sufficient and Fe-deficient medium. Means within each condition with the same letter are not significantly different according to one-way ANOVA followed by post hoc Tukey test, P < 0.05.

To confirm that the suppression of *hy5* mutant phenotype in the double mutant is because of lack of *bts* expression in the *hy5/bts* double mutant, we generated three independent BTS overexpression lines under *hy5* background (P6, P1 and P3) by transforming *Pro35S:BTS* into the *hy5* mutants. Next, we confirmed the *BTS* expression in the overexpression lines by performing qRT-PCR and found that P1 and P3 showed higher expression level. (Figure S6). We further assessed the phenotype of *Pro35S:BTS/hy5* plants using two independent T3 lines (P1 and P3) and found that the *Pro35S:BTS/hy5* plants clearly showed phenotype similar to *hy5* under -Fe (Figure 7C, D). Taken together, these results confirm that HY5 functions upstream to *BTS*.

### *HY5* is required for inducing the expression of Fe-mobilizing coumarin biosynthesis genes under Fe deficiency

*HY5* is known to induce the expression of flavonoid biosynthesis genes and promote flavonoid accumulation (Oyama, Shimura and Okada, 1997a; Holm *et al*., 2002; Shin, Park and Choi, 2007; Young *et al*., 2008; Stracke *et al*., 2010). Both flavonoids and coumarins are synthesised through the phenylpropanoid pathway (Vogt, 2010). The coumarin production and secretion increases under Fe deficiency. Since FCR activity and the expression of Fe uptake genes was compromised in the *hy5* mutant under Fe deficiency, we also analysed the expression of genes involved in coumarin biosynthesis and secretion. Our data indicates that induction of *F6’H1* (the gene which is involved in the first step of coumarin biosynthesis, (Rodríguez-Celma *et al*., 2013; Schmid *et al*., 2014)expression in response to Fe deficiency was significantly reduced in the *hy5* mutant as compared to the WT plants (Figure 8A). The same pattern was observed for *S8H* and *CYP82C4* which play a key role in the biosynthesis of fraxetin and sideretin (the main Fe-mobilizing coumarins) respectively (Figure 8A) (Rajniak *et al*., 2018; Siwinska *et al*., 2018; Tsai *et al*., 2018). We next analysed the expression of *BGLU42* (β-glucosidase) and *PDR9* (ABCG transporter) which are involved in the secretion of coumarins in the rhizosphere (Fourcroy *et al*., 2014; Zamioudis, Hanson and Pieterse, 2014). *BGLU42* expression was reduced in the *hy5* mutant under Fe deficiency but *PDR9* expression was not affected significantly in the *hy5* mutant as compared to the wild type (Figure 8B). The expression levels of *MYB10* and *MYB72* which regulate the expression of some of the coumarin biosynthesis and secretion genes, (Palmer *et al*., 2013; Zamioudis, Hanson and Pieterse, 2014) was not affected in the *hy5* mutant as compared to the WT (Figure 8C). These results indicate that HY5 function is crucial to induce the expression of Fe-mobilizing coumarins under Fe deficiency.

**Figure 8.**
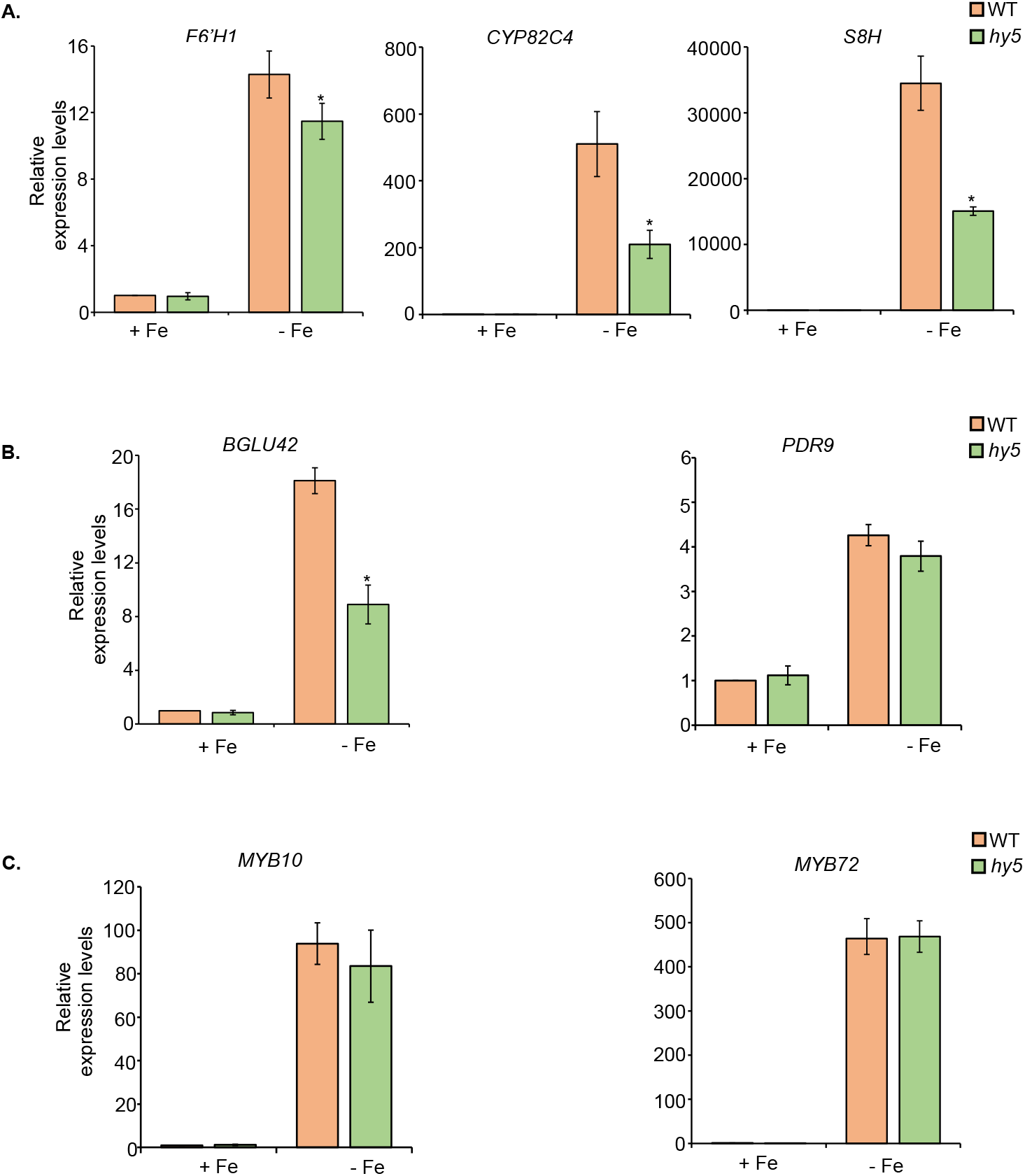
Biosynthesis of Fe-mobilizing coumarins is affected in the *hy5* mutant. A. Expression levels of *F6’H1, CYP82C4* and *S8H* involved in Coumarin biosynthesis. B. Expression levels of *BGLU42* and *PDR9* involved in coumarin secretion. C. Expression levels of *MYB10* and *MYB72* genes involved in transcriptional control of coumarin biosynthesis and secretion. Relative expression was determined by qRT-PCR in WT and *hy5* mutant seedlings grown on +Fe media for 6 days and transferred to both +Fe and –Fe (+300µM Fz) for three days. Data shown is an average of three biological replicates (n=2 technical replicates). Each biological replicate consists of pooled RNA extracted from roots of ∼ 90 seedlings. Error bars represent ±SEM.*Significant difference by Student’s t test (P≤0.05).

Coumarins are known to fluoresce when exposed to 365nm UV light (Dorey *et al*., 1997; Ahn *et al*., 2010). We analysed the fluorescence in the roots of *hy5* mutants and compared with the WT under both +Fe and -Fe conditions. In WT seedlings, fluorescence increased under -Fe conditions but in the case of *hy5* mutants fluorescence was reduced under -Fe conditions which is in agreement with the qRT-PCR results and clearly suggest that root secreted fluorescent coumarins was reduced in the *hy5* mutants as compared to WT under -Fe conditions. (Figure S7).

## Discussion

Plants utilize transcriptome reprogramming to maintain Fe homeostasis under varying Fe conditions. Several of the bHLH family TFs play an important role in regulating the expression of genes involved in maintaining Fe homeostasis (Hindt and Guerinot, 2012). Very recently, it has been shown that HY5 regulates Fe uptake in tomato (Guo *et al*., 2021). Both the leaves and roots accumulate HY5 protein in a phyB-dependent manner under Fe deficiency. *HY5* activates the expression of *FER* which leads to upregulation of transcripts involved in Fe uptake (Guo *et al*., 2021). In *Arabidopsis, HY5* is known to regulate expression of genes involved in sulphur, nitrogen and copper uptake. It regulates sulfur uptake by activating the expression of *APR1, APR2* (ADENOSINE 5’-PHOSPHOSULFATE REDUCTASE) and SULFATE TRANSPORTER 1;2 (*SULTR1;2*) involved in sulfur uptake (Lee, Koprivova and Kopriva, 2011). *HY5* along with *HYH* promotes nitrogen signalling by positively regulating NITRATE REDUCTASE 2 (*NIA2*) and NITRITE REDUCTASE 1 (*NIR1*) and negatively regulates nitrate uptake genes NITRATE TRANSPORTER 1.1 (*NRT1*.*1*) and AMMONIUM TRANSPORTER 1;2 (*AMT1;2*). *HY5* regulates copper signalling through SQUAMOSA PROMOTER BINDING PROTEIN-LIKE 7 (*SPL7*) which regulates the expression of copper uptake genes (Zhang *et al*., 2014). However, it remains unclear whether *HY5* plays any role in regulation of Fe homeostasis in *Arabidopsis*.

In our study, we characterized the role of *HY5* in regulating Fe deficiency response in Arabidopsis roots. Our phenotypic analysis revealed that *hy5* mutants are more sensitive to -Fe conditions. We found that under -Fe conditions, the overall growth of the *hy5* mutants was strongly inhibited. We compared the primary root length and chlorophyll content of the *hy5* mutants with the wild type plants under both +Fe and -Fe conditions. Our results revealed that *hy5* mutants have significantly shorter roots and reduced chlorophyll content as compared to the wild type plants under -Fe conditions (Figure 1B). Furthermore, the root length phenotype was partially rescued in the complementation lines *ProHY5:HY5:eGFP/hy5* (P-13 and P-8) (Figure 1B). The percentage reduction in the chlorophyll content under -Fe was less in the complementation lines *ProHY5:HY5:eGFP/hy5* (P-13 and P-8) as compared to the *hy5* mutants. In the alkaline soil also, we found that *hy5* mutants were more chlorotic and the chlorotic phenotype was partially rescued in the complementation *ProHY5:HY5:eGFP/hy5* (P-13 and P-8) lines. Moreover, we found that the chlorotic phenotype of the *hy5* mutants was recovered when exogeneous Fe was supplied (Figure 2A). Taken together, these results indicate that HY5 function is crucial for primary root growth and chlorophyll synthesis under -Fe conditions.

The iron content analysis by Perls staining revealed that *hy5* mutants have less iron as compared to the wild type plants and it was found to be recovered in the complementation line *ProHY5:HY5:eGFP/hy5* (P-13) (Figure 2B). This indicates that *HY5* is essential for iron accumulation in roots and this reduced accumulation of Fe in *hy5* mutants might explain the growth defects observed under Fe deficiency.

*HYH* (*HY5* homolog) is known to play overlapping roles and act redundantly with *HY5* in the regulation of hypocotyl and lateral root growth (Holm *et al*., 2002; Gangappa and Botto, 2016). In nitrogen signalling, *HYH* is known to act with *HY5* in positively regulating the expression of nitrate reductase involved in nitrogen assimilation (Jonassen, Lea and Lillo, 2008). Based on our phenotyping results, we found that *hyh* mutants showed phenotype similar to the WT plants under -Fe. In the *hy5* mutants, the root growth was inhibited under -Fe conditions but in the *hyh* mutants root length increased like WT plants (Figure S2). These results indicate that *HY5* functions independently of *HYH* in regulation of Fe deficiency responses in *Arabidopsis* (Figure S2).

The Fe deficiency responses are governed by a complex regulatory network. FIT is one of the crucial regulators of the Fe deficiency responses. It is upregulated at transcriptional level under Fe deficiency. It interacts with bHLH Ib TFs to trigger the expression of Fe-uptake associated genes (i.e. *IRT1* and *FRO2*). Our qRT-PCR analysis revealed that *HY5* positively regulates the expression of *FIT* as well as *IRT1* and *FRO2*. (Figure 3A). To independently confirm this *in planta*, we used *IRT1* transcriptional reporter line and we found that the *IRT1* induction was significantly impaired in the *hy5* mutant as compared to the WT (Figure 4). The FCR activity was also found to be less induced in the *hy5* mutant as compared to the WT under -Fe conditions (Figure S4). These results further support our qRT-PCR results that *HY5* positively regulates the activation of Fe-uptake genes under -Fe.

Fe homeostasis in plants is tightly regulated to supply required amounts of this element for an optimal growth while avoiding excess accumulation to prevent oxidative stress. Transcriptional regulatory cascade plays important role controlling Fe homeostasis and the expression of genes involved in Fe uptake is induced under Fe deficiency and turned off under Fe excess conditions. BTS, an E3 ubiquitin ligase plays an important role in maintaining Fe homeostasis by promoting degradation of bHLH105 and bHLH115 under Fe sufficient conditions, bHLH105 and bHLH115 positively regulate the expression of Fe uptake genes (Selote *et al*., 2015). *BTSL1* and *BTSL2* are two *BTS* paralogs and they also negatively regulate Fe uptake by targeting FIT for degradation (Hindt *et al*., 2017; Rodríguez-Celma *et al*., 2019). The regulation of *BTS* expression is important to avoid Fe overload. In our study, we found that HY5 binds to the *BTS* promoter and negatively regulates *BTS* expression under Fe deficiency as our qRT and GUS staining results revealed that *BTS* is induced more in the *hy5* mutants as compared to the wild type (Figure 5A, Figure 6). The *bts* mutation in the *hy5* mutant background suppresses *hy5* mutant phenotype suggesting that *BTS* acts downstream to *HY5* (Figure 7A, B). Based on these results, we hypothesized that the *hy5* mutant phenotype under -Fe is dependent on *BTS* expression in the *hy5* mutant because in the *hy5/bts* double mutants in which *BTS* is not expressed, the *hy5* mutant phenotype is rescued. This was further supported by the *35S:BTS/hy5* (*BTS* overexpression in the *hy5* mutant) phenotyping results which revealed that *35S:BTS/hy5* also showed sensitive phenotype similar to the *hy5* mutants under -Fe (Figure 7 C, D). Altogether, these results indicate that HY5 acts upstream to *BTS* in the Fe signalling pathway and negatively regulates its expression. Similarly, we also found that HY5 negatively regulates PYE expression under -Fe (Figure 3B). Further studies are required to explore to find whether HY5 directly regulates *PYE* expression.

Coumarins have recently emerged as key players for Fe uptake particularly in high pH (Robe *et al*., 2021). They aid in the mobilisation of Fe in the soil (Tsai and Schmidt, 2017). The production as well as the secretion of coumarins into the rhizosphere increases under Fe deficiency. The expression of coumarin biosynthesis genes (i.e. *CYP82C4* and *S8H*) is activated by FIT/bHLH Ib complexes (Schmid *et al*., 2014; Tsai and Schmidt, 2017; Tsai *et al*., 2018). The MYB10, MYB72 transcription factor and FIT/bHLH Ib complexes regulate the expression of *F6’H1* involved in coumarin biosynthesis as well as *BGLU42* and *PDR9* involved in coumarin secretion (Palmer *et al*., 2013; Tsai and Schmidt, 2017). In our study, we found that *CYP82C4, S8H, F6’H1* and *BGLU42* induction was significantly lower in the *hy5* mutants as compared to the wild type under -Fe (Figure 8A, B). These results indicate that *HY5* is essential for activating the expression of genes involved in coumarin biosynthesis. In addition to this, fluorescence was found to be reduced in the *hy5* mutant as compared to the wild type under -Fe conditions which indicates that coumarin production and secretion is less induced in the *hy5* mutants as compared to the wild type (Figure S7). Altogether, these results indicate that *HY5* is essential for induction of coumarin production and secretion under -Fe.

Based on our results and existing literature, we propose a model depicting the crucial involvement of HY5 under Fe-deficiency response in *Arabidopsis thaliana* (Figure 9). HY5 positively regulates the expression of *FIT* as well as other downstream genes such as *IRT1* and *FRO2* that are involved in Fe uptake and *S8H, CYP82C4, F6’H1, BGLU42* involved in coumarin biosynthesis and secretion. Additionally, *HY5* acts as a negative regulator of *BTS and PYE*. Importantly, it directly binds to the promoter of *BTS* to regulate its expression.

**Figure 9.**
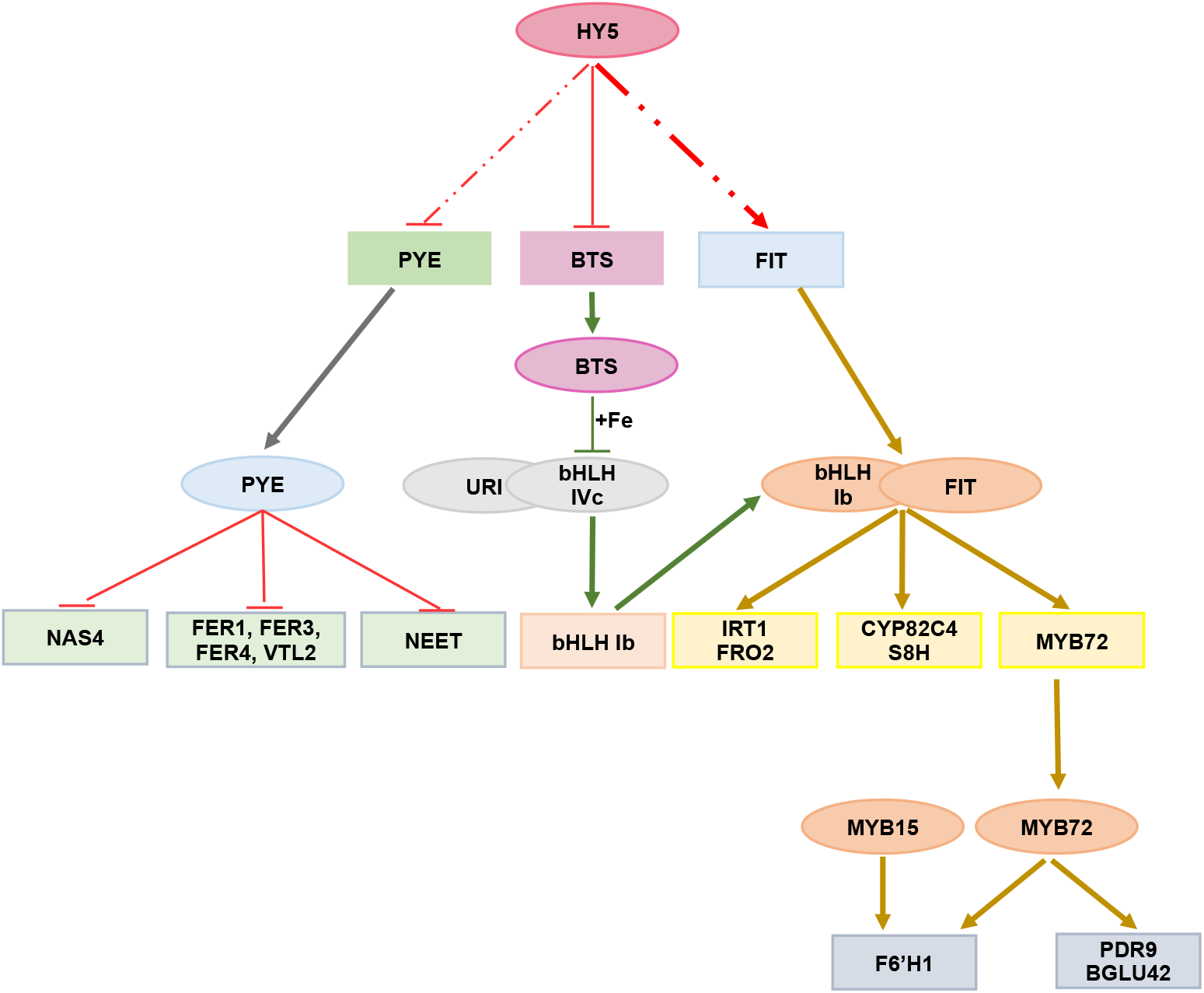
Model describing the role of HY5 in maintaining iron homeostasis. HY5 negatively regulates the expression of *BTS* and *PYE*. It directly interacts with the promoter of *BTS* and negatively regulates its expression. Under iron sufficient conditions, BTS is known to degrade bHLH IVc TFs which activate the expression of bHLH Ib TFs under iron deficient conditions. HY5 positively regulates the expression of *FIT* as well as other genes including *IRT1,FRO2* (involved in iron uptake) and *CYP82C4, S8H, BGLU42* and *F6’H1* (involved in coumarin biosynthesis) whose expression is regulated by FIT along with bHLHIb TFs. *HY5* negatively regulates PYE expression which is known to repress *NAS4*; *FER1, FER3, FER4, VTL2*; and *NEET* (involved in iron transport, storage and assimilation respectively). Oval boxes represent protein and rectangular boxes represent mRNA.

Several studies have revealed that HY5 can be both an activator and a repressor (Ang *et al*., 1998; Lee *et al*., 2007; Ruckle, DeMarco and Larkin, 2007; Kindgren *et al*., 2012; Delker *et al*., 2014; Xu *et al*., 2016; Gangappa and Botto, 2016; Norén *et al*., 2016; Zhang *et al*., 2017; Gangappa and Kumar, 2017; Nawkar *et al*., 2017; Burko *et al*., 2020; Yadukrishnan, Rahul and Datta, 2020). Our results so far clearly indicates that HY5 acts both an activator and a repressor to regulate the genes involved in Fe regulatory network.

In summary, our findings using gene expression analysis, genetic analysis and ChIP experiments provide novel evidence that HY5 regulation of *BTS* is critical for the Fe deficiency responses in Arabidopsis. Also, HY5 functions as a direct transcriptional repressor of *BTS* and potentially direct transcriptional activator of several other key genes (*IRT1, FRO2, FIT1, S8H, CYP82C4, F6’H1, and BGLU42)* involved in the Fe regulatory network. Further experiments will be required to understand how HY5 is regulated at posttranscriptional and posttranslational level under Fe deficiency and whether HY5 directly regulates the other key genes involved in Fe regulatory pathway in addition to *BTS*. Moreover, interaction studies with bHLH TFs involved in Fe signalling pathway will shed the light whether HY5 interacts with them and regulate the genes involved in Fe homeostasis. The knowledge gained on HY5 function and its role in Fe homeostasis by regulating *BTS* in Arabidopsis can be further extended to crop plants.

## Materials and Methods

### Plant Materials and Growth Conditions

In this study, Columbia (Col-0) and Wassilewskija (WS-0) ecotypes of *Arabidopsis thaliana* were used as wild type. The mutant lines used in this study were *hy5* [SALK_056405; (Saini *et al*., 2020), *bts-1* [SALK_016526; (Long *et al*., 2010)] and *pye* [SALK_021217; (Long *et al*., 2010) in the Col-0 background. In addition, following mutant lines were kindly provided by Roman Ulm: *hy5, hyh* and *hy5/hyh* in the WS background and *pHY5:HY5:YFP/hy5* in the Ler background. *hy5* single mutant was crossed with *bts-1* to generate *hy5/bts* double mutant. The *hy5* allele was also introduced into *pIRT1:GUS* line (Blum *et al*., 2014) and *pBTS:GUS* line (Selote *et al*., 2015) by crossing. The single as well as double mutants were confirmed by genotyping. The primers used for genotyping are listed in Table 1. Seeds were surface sterilized using 5% sodium hypochlorite followed by 70% ethanol and stratified in dark at 4 ºC for 3 days. Plants were grown on ½ Murashige and Skoog media (Caisson labs, USA) with Fe (+Fe) or ½ Murashige and Skoog media (Caisson labs, USA) without Fe (-Fe). The pH of media was adjusted to 5.7 with KOH. To create Fe deficiency conditions, 300µM of [3-(2-pyridyl)-5,6-diphenyl-1,2,4-triazine sulfonate] Ferrozine (Sigma-Aldrich) was added in the -Fe media. For seedling growth, plants were grown under long-day conditions, 16 h light and 8 h dark at 22ºC with 50% humidity and a light intensity of 100-110 µmol/cm^2^/s. Soil was made as a mixture of soilrite, perlite and compost (3:1:1). CaO (Sigma-Aldrich) (7.8 gm CaO/kg) was mixed in the soil to generate alkaline pH soil.

### Phenotypic Analyses

For phenotypic characterization, seeds were sown on ½ MS media with Fe (+Fe) and also on ½ MS media without Fe (-Fe). The plates were kept vertically in growth chamber and 10 day old seedlings growing on the plates were scanned using Epson Perfection V600 with 1200dpi resolution. The root length was quantified using ImageJ 1.52a software (National Institutes of Health).

### Chlorophyll Content Measurement

The chlorophyll content was measured using seedlings grown on +Fe and -Fe for 10 days. It was measured by extracting chlorophyll using 1ml of 80% acetone from leaf tissues of five to six seedlings and incubating in dark for 24 hours using the formula (mg/g) = <u>(20.3 x A</u>_<u>645</u>_ <u>+ 8.04 x A_663_) x V</u>/ W X 10^3^ (Aono *et al*., 1993).

### Ferric-Chelate Reductase Assay

Ferric-chelate reductase assay was performed as previously described with some modifications (Ying Yi and Mary Lou Guerinot, 1996). Seeds were grown for five days on +Fe and then transferred to both +Fe and -Fe (+300 µM Ferrozine) for 3 days. Roots from 30 to 40 seedlings were collected in 700 µl assay solution consisting of 0.1mM Fe (III)-EDTA and 0.3mM ferrozine in distilled water. An identical assay solution without roots was used as blank. The FCR activity is determined by taking absorbance at 562 nm with the help of a spectrophotometer using the formula (µM Fe (II)/g root FW/hr = (A/28.6) x V/ Root FW.

### Histochemical Perls Staining

For Fe staining, five days old seedlings grown on +Fe were vacuum infiltrated with a solution containing 1% (v/v) HCL and 1% (w/v) K-ferrocyanide for 5 minutes. Seedlings were then washed with water five times, observed and photographed using NIKON ECLIPSE Ni U microscope.

### RNA Extraction and qRT-PCR

Total RNA was extracted from roots of seedlings grown on ½MS for six days and transferred to +Fe or -Fe for three days using Plant RNeasy kit (Qiagen) following manufacturer’s protocol and digested with DNase I to remove genomic DNA. cDNA was synthesised using RevertAidTM First Strand cDNA Synthesis Kit (Thermo) from RNA (2µg). qPCR was performed using a LightCycler 480 II (Roche) and TB GreenTM Premix Ex. The β-tubulin was used as a reference gene and relative expression levels were calculated using comparative threshold cycle method *(ΔΔCT*). The primers used are listed in Table 2.

### Histochemical GUS Staining

GUS staining was done using GUS-expressing seedlings grown on +Fe or -Fe for 8 days. For visualization of GUS expression, seedlings were incubated in staining solution (100 mM Na_2_HPO_4_, 100 mM NaH_2_PO_4_, 2 mM K_4_[Fe(CN)_6_], 2 mM K_3_[Fe(CN)_6_], 0.2 Triton X-100, and 2 mM 5-bromo-4-chloro-3-indolyl glucuronide, pH 7.0) and vacuum was applied for 25 minutes followed by incubation at 37°C for 2 to 3 hours. After incubation, the seedlings were stored in 70% ethanol before imaging. Images were taken using NIKON ECLIPSE Ni U microscope.

### Confocal Microscopy

For confocal microscopy, seedlings grown on ½ MS for seven days were stained with 10 μM propidium iodide for 45 seconds. For GFP excitation, a 488nm laser was used and PI was excited with 561nm laser and emission spectra were collected at 500-530 nm and 600-650nm respectively. Images were taken using a SP8 upright confocal microscope (Leica).

### Fluorescence

Seeds were sterilized and sown on +Fe and -Fe for 10 days. Fluorescence of 10-d-old plants was observed using a Gel Doc XR+ System (Biorad) with an excitation wavelength of 365nm (Vanholme *et al*., 2019).

### Plasmid constructs and transgenic lines

For overexpression lines, the coding sequence of *BTS* was amplified from cDNA. The forward primer was designed with a CACC overhang to facilitate directional cloning of the CDS fragment into pENTR/D-TOPO. The resulting clone was sequence verified and used to set up an LR reaction with the destination vector (pMDC32) to generate *35S:BTS*. The construct was introduced into *Agrobacterium tumefaciens GV3101* and transformed into wild-type Col-0 background by the floral dip method. Single gene insertions were detected by T2 3:1 segregation ratio on media containing hygromycin followed by homozygous line selection at T3.

For *pHY5:HY5:eGFP* construct, the *HY5* gene was amplified from cDNA and cloned in pENTR/D-TOPO. The resulting clone was digested with PspOMI and StuI and the *HY5* coding sequence was cloned at the N terminus of eGFP in pENTR/D/TOPO eGFP vector. The native promoter (750 bp upstream of the translational start codon) was amplified from gDNA and cloned into SacI and KpnI restriction sites in pAtExp7 containing attR1 and attR2 recombination sites. The resulting clone was used to set up a LR reaction with pENTR/D/TOPO HY5-eGFP to create *pHY5:HY5-eGFP* which was then transformed into *hy5* mutant. The primers used are listed in Table 3.

### ChIP-qPCR

ChIP was performed according to the method described previously (Gendrel *et al*., 2005). *pHY5:HY5:YFP/ hy5* and *35S:eGFP* in the wild type Ler background grown and germinated on ½ MS media for 7 days and then transferred to ½ +Fe and ½ -Fe for 3 days were fixed using 1% formaldehyde. Nuclei isolation was done followed by shearing using a Qsonica 800R ultrasound sonicator with 30 cycles of 70% pulse amplitude for 15s followed by 45s pulse off time. The protein-A magnetic beads (1614013, SureBeads) and anti-GFP antibody (a290, Abcam) were used to pull down HY5-DNA complexes. Rabbit IgG was used as a negative control. The immunocomplexes bound to beads were washed and eluted from beads followed by reverse crosslinking. The qPCR was set up using precipitated chromatin. The Fold enrichment was determined by normalizing against negative control (IgG). The primers used are listed in Table 4.

### Statistical Analysis

Data was expressed as means ± SEM. P-value was calculated by student’s t test or one-way ANOVA followed by post-hoc Tukey HSD Test. P-value ≤ 0.05 was considered as statistically significant.

## Acknowledgments

We thank Roman Ulm for providing WS, *hyh, hy5, hyh/hy5* and *pHY5:HY5:YFP/hy5* (Ler) seeds. We are grateful to Rumen Ivanov and Petra Bauer for providing us the *pIRT1:GUS* seeds. Terry A. Long for *pye, bts* and *pBTS:GUS* and Janneke Balk for *bts* seeds. We thank Prince Saini and Asis Kumar for their technical help in qRT-PCR and ChIP-qPCR experiments. We are grateful to members of SBS laboratory, Ramesh Yelagandula, Balaji Enugutti, Kaushal Bhati and Marco Giovannetti for critical reading of the manuscript. S.B.S. acknowledges intramural funding support from Indian Institute of Science Education and Research (IISER) Mohali. S.B.S. is a recipient of Ramalingaswami Fellowship from Department of Biotechnology (DBT) Govt. of India and acknowledges funding received from DBT. SBS also acknowledges Science and Engineering Research Board (SERB) for Early career research funding (ECR/2018/001580). S.M. and D.S. acknowledges PhD fellowship and postdoctoral fellowship received from IISER Mohali.

## Author contributions

S.M. and S.B.S. conceived the study and designed the experiments. SM performed all the experiments with the help from D.S., K.M., H.M., and V.M. S.M., S.B.S., R.K.Y. and A.K.P. analysed the data. S.M. prepared the figures and wrote the first draft. S.B.S. supervised the work, edited the manuscript, and provided the overall direction. All authors discussed the results and commented on the manuscript.

