## Supplementary data for "Elongated Hypocotyl 5 (HY5) regulates *BRUTUS* (*BTS*) to maintain Iron homeostasis in *Arabidopsis thaliana*"

*pHY5:HY5:eGFP/hy5 (P-13)* *pHY5:HY5:eGFP/hy5 (P-8)*

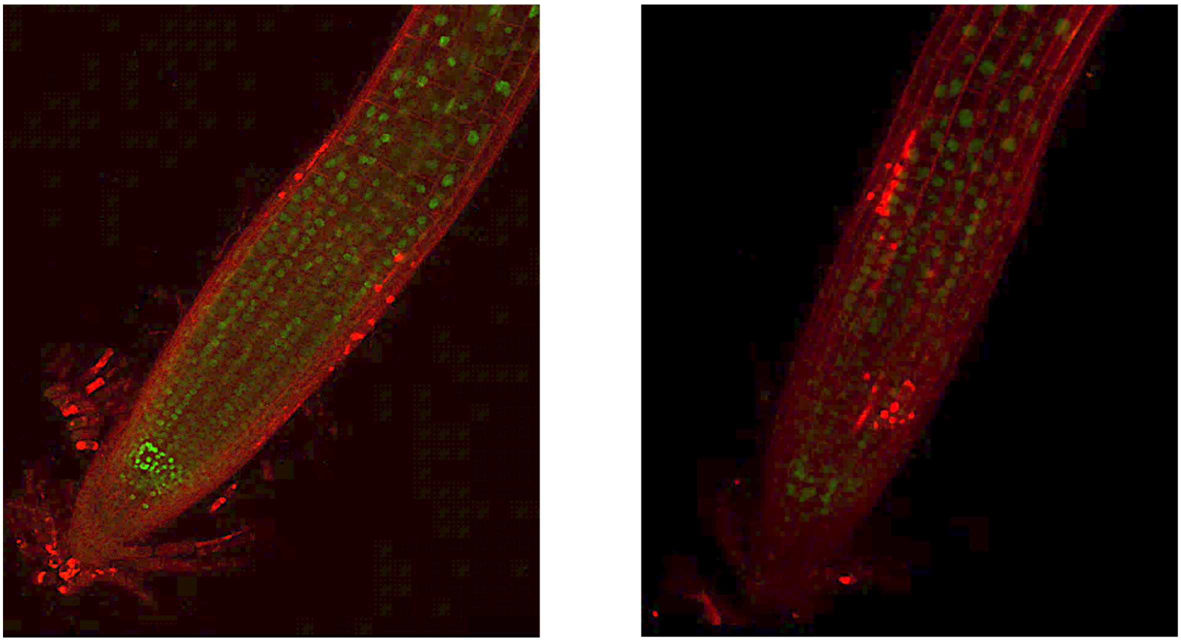

**Figure S1.** HY5 is expressed in all the cell layers in root.
Confocal microscopy of roots from the complementation lines *pHY5:HY5:eGFP/hy5 (P-13 and P-8).* Seedlings grown on +Fe media for 7 days were stained with PI for imaging.

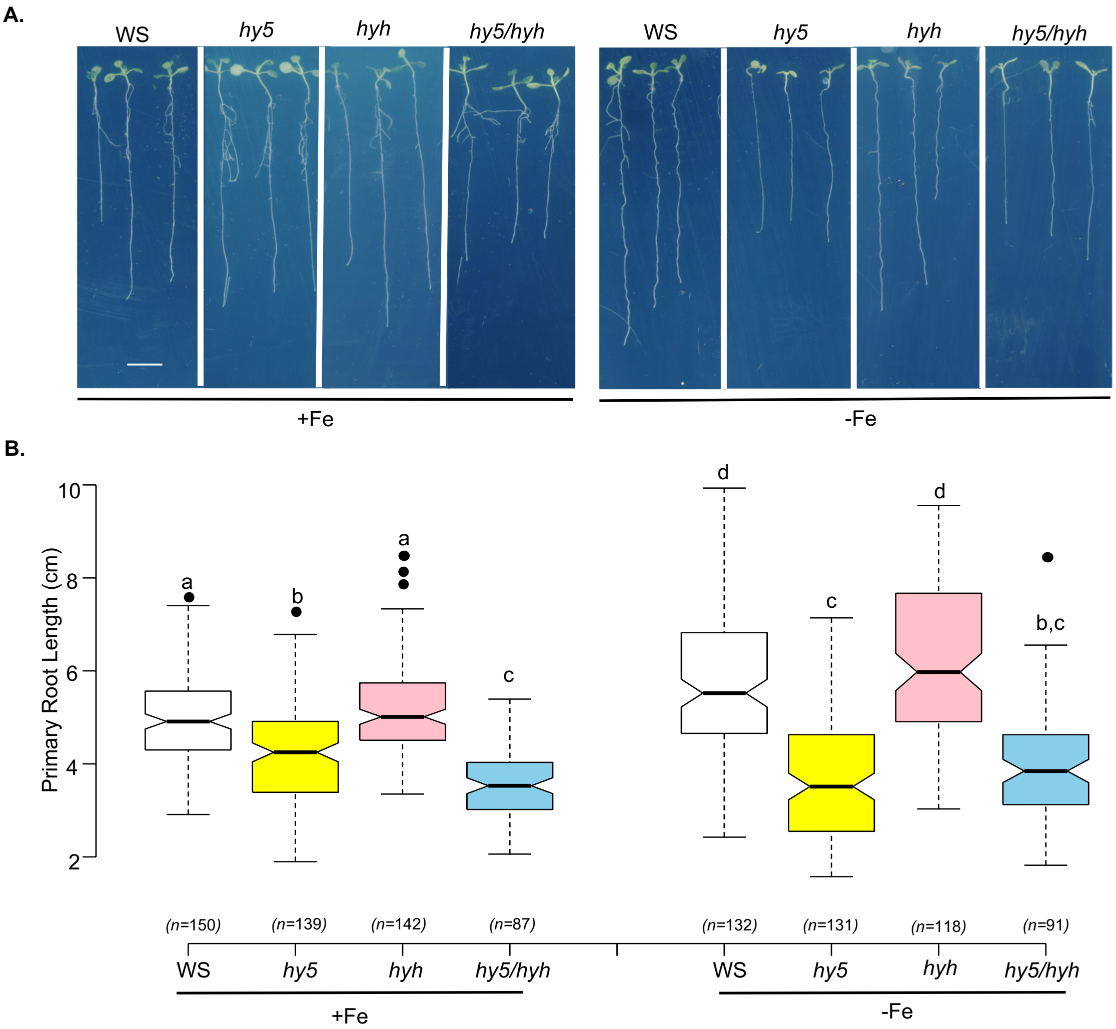


**Figure S2.** HY5 regulates primary root growth under iron limiting conditions independently of HYH.

**A**. Phenotypes of the *Arabidopsis* wild type (WS), *hy5*, *hyh* and *hy5hyh* grown for 10 days on Fe-sufficient and Fe-deficient medium. Scale bars, 1cm. **B.** Boxplot of **r**oot length of the wild type (WS), *hy5*, *hyh* and *hy5hyh* grown for 10 days on Fe-sufficient and Fe-deficient medium. Means within each condition with the same letter are not significantly different according to one-way ANOVA followed by post hoc Tukey test, $P < 0.05$.

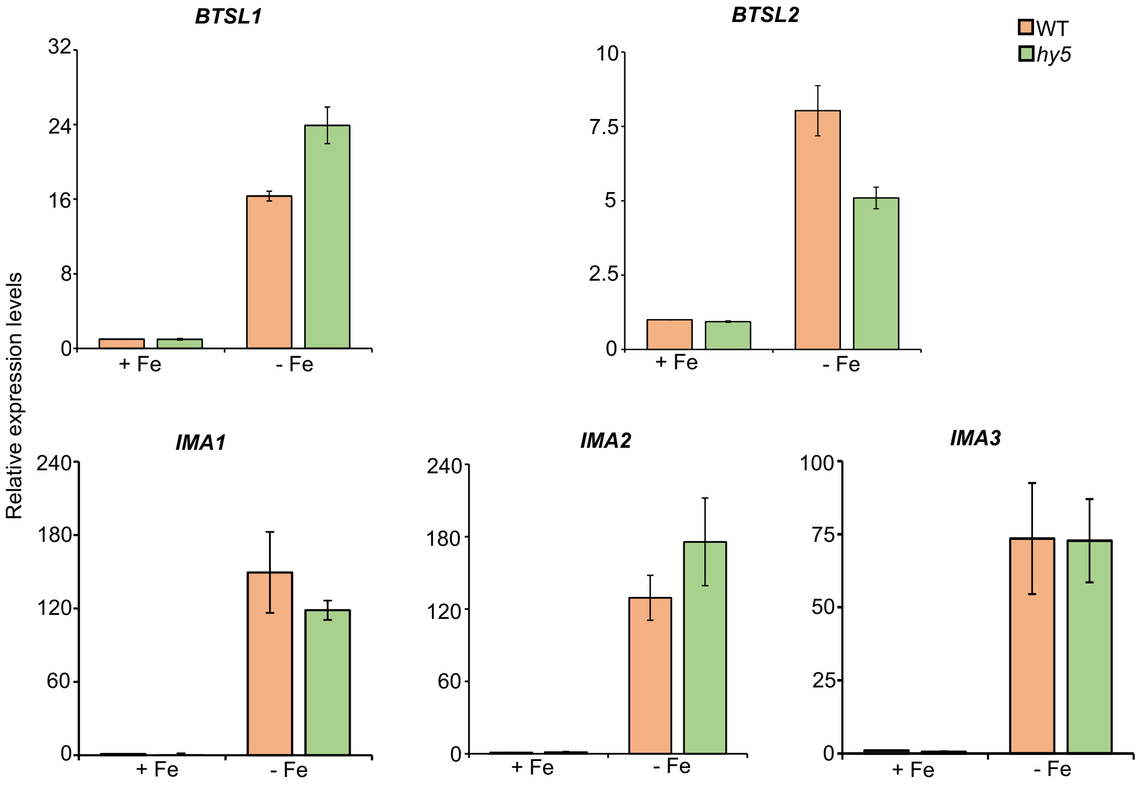


**Figure S3.** Expression levels of genes involved in iron deficiency in the *hy5* mutant.
Expression levels of *BTSL1*, *BTSL2, IMA1, IMA2* and *IMA3*. Relative expression was determined by qRT-PCR in WT and *hy5* mutant seedlings grown on +Fe media for 6 days and transferred to both +Fe and –Fe (+300µM Fz) for three days. Data shown is an average of three biological replicates. Error bars represent ±SEM.*Significant difference by Student's t test (P≤0.05).

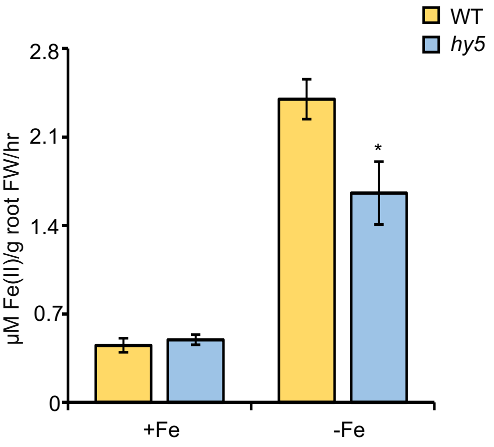


**Figure S4.** Ferric-chelate reductase activity of the wild type (WT), *hy5* and grown for 5 days on +Fe and transferred to +Fe-sufficient or –Fe (+300µM Fz) for 3 days. Data shown is an average of three independent experiments. Each experiment consists of five to six biological replicates and each biological replicate consists of a pool of around 50 roots. Error bars represent ±SEM. Significantly different (*) according to one-way ANOVA followed by post hoc Tukey test, $P < 0.05$.

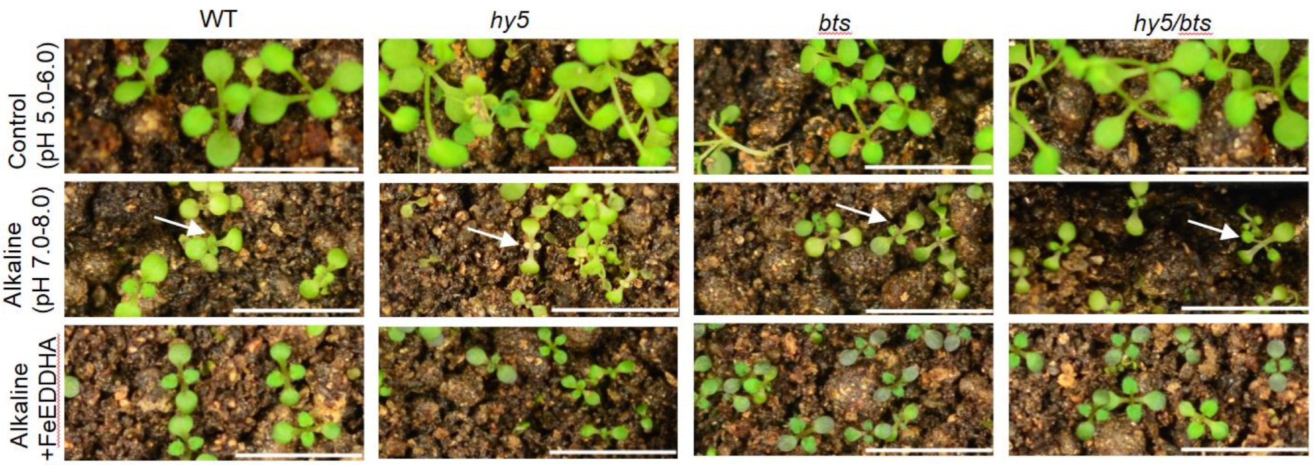


**Figure S5.** *bts* mutation suppresses the *hy5* mutant phenotype. Phenotypes of the WT, *hy5, bts* and *hy5bts* grown on normal soil (pH 5.0-6.0), alkaline soil (pH 7.0-8.0) and alkaline soil watered with Fe-EDDHA (Ethylenediamine di-2-hydroxyphenyl acetate ferric) for two weeks. Bars =10mm. White arrows indicate true leaves undergoing chlorosis.

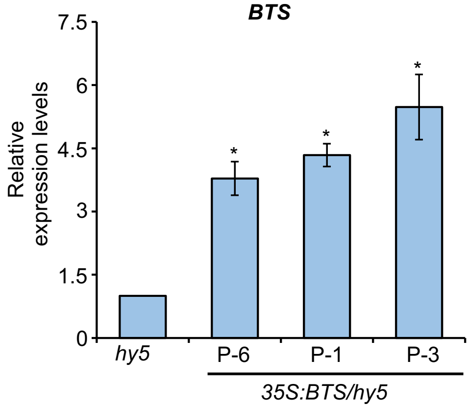


**Figure S6.** Overexpression of *BTS* in the *hy5* mutant.
qRT-PCR analysis of *BTS* expression in *hy5* mutant and BTS overexpression lines under *hy5* mutant background (P-6, P-1 and P-3). Relative expression was determined by qRT-PCR in *hy5* and *35S:BTS/hy5* (P-6, P-1 and P-3) seedlings grown on +Fe media for one week. Data shown is an average of two biological replicates (n=2 technical replicates). Each biological replicate consists of pooled RNA extracted from ~ 90 seedlings. Error bars represent ±SEM.*Significant difference by Student's t test (P≤0.05).

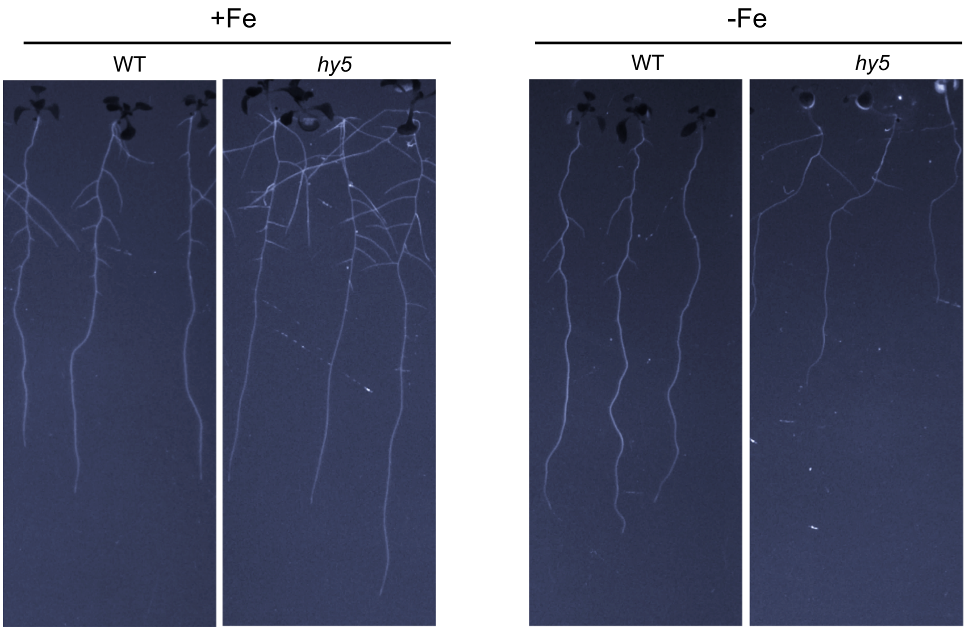


**Figure 7.** Roots of *hy5* mutants show reduced fluorescence under -Fe. Visualization of fluorescent phenolic compounds produced and secreted by the roots of 10 day old WT and *hy5* seedlings grown for 10 days on +Fe and –Fe. The contrast/brightness was adjusted for good visualization.

Table 1. Primers used in genotyping.

| | |
|---|---|
| bts-1, LP | CCAAATGCGTTCGTAGGTAAG |
| bts-1, RP | TCAGATTTACACAAATTTGCAGC |
| hy5, LP | TTCACTCTCGATATCCGTTCG |
| hy5, RP | ATGCGAGTGAATGACCATTTC |
| LBb1.3 | ATTTTGCCGATTTCGGAAC |

Table 2. Primers used for qRT-PCR.

| | |
|---|---|
| qbHLH38 F | ACGGTGCCGGAGATAACCTA |
| qbHLH38 R | GTCGGTCACGTTCACTAGCA |
| qbHLH39 F | CCGTTCATGTCTTCCTGCCT |
| qbHLH39 R | GCCTTTGGTGGCTGCTTAAC |
| qbHLH100 F | CTCCCACCAATCAAACGAAGAAG |
| qbHLH100 R | TGTTTTGGTCGGTGTAAACGAG |
| qbHLH101 F | AAGAAGATCGAGGAGCGGTG |
| qbHLH101 R | TGTTTTGGTCGGTGTAAACGAG |
| qRT PYE FP | CAGGACTTCCCATTTTCCAA |
| qRT PYE RP | CTTGTGTCTGGGGATCAGGT |
| qRT FRO2 FP | GCCACATCTGCGTATCAAGTT |
| qRT FRO2 RP | TCCCAAACAAGCTACGACCA |
| qRT FIT FP | CAGTCACAAGCGAAGAAACTCA |
| qRT FIT RP | CTTGTAAAGAGATGGAGCAACACC |
| qRT BTS FP | GCTCTGGCACAAGTCAATCA |
| qRT BTS RP | CGTTCATCAAATGCCGATAA |
| qRT IRT1 FP | GAATGTGGAAGCGAGTCAGCGA |
| qRT IRT1 RP | GATCCCGGAGGCGAAACACTTA |
| qRT TUBULIN FP | CGACAATGAAGCTCTCTACGA |
| qRT TUBULIN RP | AAGTCACACCGCTCATTGTT |
| qRT MYB10 FP | GGGGAAATCTTGGTGGAGCA |
| qRT MYB10 RP | AGGAGGAACCTGGCTATCGT |
| qRT MYB72 FP | TCGAGAGGTAACCAAATCGCA |
| qRT MYB72 RP | CAGCTGTCTCCTCAAGTCGG |
| qRT NAS4 FP | GGCTTCGACGTTGTGTTCTT |
| qRT NAS4 RP | AGCAAAGCACCAGGAGACAT |
| qRT At-NEET FP | TCGTTGTCACCGAGCTTTCC |
| qRT At-NEET RP | ACGTCCCCGACCTCCAA |
| qRT F6'H1 FP | TGATATCTGCAGGAATGAAACG |
| qRT F6'H1 RP | GGGTAGTAGTTAAGGTTGACTC |
| qRT S8H FP | CCGAGACACTTGGCTTCTT |
| qRT S8H RP | CAGCAGCTCCACCGAAACA |
| qRT CYP82C4 FP | AGGCTCAGTATCGTCGGAG |
| qRT CYP82C4 RP | TTTCTATGTCTGAATCCTCGACG |
| qRT PDR9 FP | GTCTTGGACACTCAACGGGT |
| qRT PDR9 RP | ATCTTGCAACCGTCGTGGAT |
| qRT BGLU42 FP | ATGGCCTGGGAACTGAAGTC |
| qRT BGLU42 RP | ATTTGTCCAACCTCCGATTG |
| qRT IMA1 FP | ATGTCTTTTGTCGCAAACTT |
| qRT IMA1 RP | CACCACCATTCTCACTATATG |
| qRT IMA2 FP | TGCTTCCGTGGTGTATGTTG |
| qRT IMA2 RP | CAAGAAAACTCGAGACACAATC |
| qRT IMA3 FP | GGCAGGCTATACGAATCAAC |
| qRT IMA3 RP | GTTCTATGTCAAGAAGCACA |
| qRT BTSL1 FP | GGCAATGAAGATGGATTTGG |
| qRT BTSL1 RP | TCATATGGAACCGTTGCTGA |
| qRT BTSL2 FP | CGGGGCAGAATCCATCTTAT |
| qRT BTSL2 RP | GTTGCAACAAGGAGCAAGAAG |

Table 3. Primers used in Cloning.

| | |
|---|---|
| pHY5 FP/Sac1 | CACCGAGCTCTCTAATGTTAACGTTGAGATGG |
| pHY5 RP/Kpn1 | AAGGTACCTTTTCTTACTCTTTGAAGATCG |
| HY5 CDS FP | CACCATGCAGGAACAAGCGACTAGCTC |
| HY5 CDS RP/ StuI* | TAAAAGGCCTAAGGCTTGCATCAGCATTAGAACC |
| BTS CDS, FP/SpeI | CACCACTAGTATGGCGACGCCGTTACCAGAT |
| BTS CDS,RP/StuI-STOP | ATAATAGGCCTTCAGGATGAGGTTGAGCAGTCC |

Table 4. Primers used for ChIP qPCR.

| | |
|---|---|
| pBTS ChiP FP | CTCCTTCTAACTCCGAGAAC |
| pBTS ChiP RP | GAAAATGAATAAAAGTGCTTGG |
| pBTSChiP ii FP | GAAAAAAAAGGAATGTGTTG |
| pBTSChiP ii RP | TTGATGAAGTAGAAAATGGTG |
